# Accurate determination of the preferred aggregation number of a micelle-encapsulated membrane protein dimer

**DOI:** 10.1101/2025.10.02.679921

**Authors:** Jonathan J. Harris, George A. Pantelopulos, John E. Straub

## Abstract

The preferred aggregation number of dodeclyphoshocholine (DPC) micelles 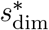 encapsulating dimeric and higher order protein assemblies is difficult to determine via experimental techniques due to uncertainty in dimer geometry and heterogeneity in the conformational ensemble. Dimerization of the Amyloid Precursor Protein transmembrane domain (C99) is a particular step of importance in the production of amyloid-*β* protein and the amyloid cascade. Molecular dynamics simulations of the C99 dimer and other transmembrane proteins have been performed to compliment micelle-phase protein structure studies. It has often been assumed that the value of 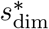 is the same as that of the pure, empty micelle. Here, we provide a convenient method for testing that assumption, while also accounting for the finite-size effects inherent in computer simulations of micelle self-assembly. Employing large, unbiased, coarsegrained molecular dynamics simulations of DPC and C99 dimer self-assembly, we determined the radius of gyration to be 21.6 ± 2.0 Å for the micelle-encapsulated dimer, and 16.0 ± 1.0 Å for the pure DPC micelle. Using these radii of gyration, we performed all-atom simulations of DPC-encapsulated C99 dimers with preferred aggregation numbers of 100 and 54 DPC to test the effect of using an expected versus a naive estimate of aggregation number on the structure of the transmembrane protein dimer. Through atomistic simulations, we determined that the transmembrane dimeric structure displays different characteristics depending on the aggregation number of the micelle, in addition to increased water penetration and micelle defects when the aggregation number is too small.

## Introduction

Surfactant micelles are significant for their biophysical utility as membrane mimics. In particular, dodecylphosphocholine (DPC) micelles have been used in conjunction with nuclear magnetic resonance (NMR) for determining the transmembrane structure of the C99 homodimer.^1^ C99 is the 99-residue C-terminal domain of Amyloid Precursor Protein, cleaved by *γ*-secretase at several sites to produce amyloid-*β* peptide, which is on-pathway for the amyloid cascade long-associated with with Alzheimer’s disease.^2–6^ Accurate modeling of the C99 dimeric structure is essential for elucidating the pathology of this protein fragment and informing future drug discovery work.

The structure of the DPC micelle-encapsulated C99 dimer has been characterized by the molecular dynamics (MD) simulations of Dominguez et al.^7^ These simulations provided a quantitative comparison between the structure of the dimer in a DPC micelle with that of the dimer embedded in a planar lipid bilayer. There were significant differences in dimeric structures between the two hydrophobic environments, notably that the dimer formed a crossed “X” conformation observed in micelle, and a parallel “||” alignment in the bilayer. Similarly, the structure of the C99 monomer was shown to depend membrane curvature, through comparison of MD simulations of the peptide inside membranes and micelles.^8^ These findings taken together prompt further investigation into the effect of micelle size on the protein’s structure, and, even more fundamentally, the expected aggregation number of a given surfactant around the protein. It is not clear, for example, how many DPC surfactants are involved in encapsulating the C99 homodimer in the NMR experiments used to determine the structure.^1^ In the work of Dominguez et al. for the C99 homodimer, the preferred aggregation number was assumed to be the same as the established experimental aggregation number for pure DPC micelles, around 54.^9,10^ The micelle-encapsulated dimer was then constructed pre-formed based on this number of surfactants as the initial structure for MD simulations.

This assumption about the size and structure of pre-formed micelle-encapsulated proteins has been investigated before, notably by Bond et al.^11^ for DPC surfactants with OmpA and Glycophorin A (GpA), and Braun et al.^12^ for SDS surfactants with the GpA dimer. These studies both concluded unsurprisingly that the protein structure will be the same regardless of whether the micelle was pre-formed or self-assembled. The crucial shortcoming of these studies is that they neglect finite-size effects inherent in computer simulations of micelle self-assembly.^13–15^ Modeling self-assembly around a single protein only allows the formation of a single micelle-encapsulated protein, which will introduce a significant bias for the resulting preferred aggregation number. Based on our previous work on the self-assembly of pure DPC micelles with coarse-grained (CG) MD simulations,^15^ it is expected that sufficient surfactants for the formation of at least three mi-celles of the correct preferred aggregation number are necessary to avoid finite-size effects. Finite-size effects for self-assembly around a protein are slightly more complicated, considering that there should be sufficient surfactants for the formation of both micelle-encapsulated proteins and pure (or “empty”) micelles. A similar analysis of the assembly of nanodiscs in the presence of peptides was conducted by Pastor and coworkers with a highly coarse-grained model.^16^

To compute the most accurate preferred aggregation number of the DPC micelle-encapsulated C99 dimer, we constructed large systems represented by a CG force field, MARTINI2.2,^17^ with 5 C99 dimers and 500 DPC surfactants. An important finding in our previous MARTINI2 micelle study^15^ is that, while the CG model yields aggregation numbers that are significantly smaller than experimental values, a micelle built with an atomistic force field with the same radius of gyration (*R*_*g*_) does match the experimental aggregation number. We exploit this observation in the current study to build pre-assembled atomistic DPC micelle-encapsulated C99 dimers based on the preferred *R*_*g*_ established by the large CG simulations. The dimer can then be simulated inside a micelle with an accurate preferred aggregation number and analyzed in atomistic detail.

The the paper is organized as follows. The next section details the methods for the CG simulations of micelle self-assembly in the presence of 5 C99 dimers and 500 DPC surfactants, followed by the description of the atomistic micelle-encapsulated C99 dimer simulations for micelles with *s*^∗^ = 100 and *s*^∗^ = 54 surfactants. The results section first provides a thorough analysis of the CG simulations, highlighting satisfactory distributions of preferred aggregation numbers for both dimer and empty micelles, and presenting the preferred *R*_*g*_ for this system. The second part of the results section is a comparison of dimeric and micellar structure between the two aggregation numbers.

## Methods

### Determination of preferred aggregation number of micelle-encapsulated C99 dimer with MARTINI simulations

Large systems of surfactants and C99 dimers were simulated from random initial configurations in order to best capture the thermodynamic preferred aggregation number of the C99 dimer (which will be referred to as 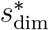) without suffering from the inaccuracy of finite-size effects. Only the C99 transmembrane and juxtamembrane domains (residues 16-55) are modeled, as in past experimental and MD work focused on characterizing the transmembrane domain.^2^ The systems consisted of 5 C99 dimers, 500 dodecylphosphocholine (DPC), 91564 water beads (of which 9156 were antifreeze waters), 52 sodium ions, and 82 chloride ions represented by the MARTINI v2.0 coarse-grained (CG) force field.^17^ The formation of 5 independent micelle-encapsulated dimers with an excess of surfactants was deemed to be adequate to approximate the thermodynamic limit of micellization based on our previous work.^15^

In order to determine the time needed for sufficient equilibration of the CG micelle system, we invoked the ergodic measure of Thirumalai et al.,^18,19^

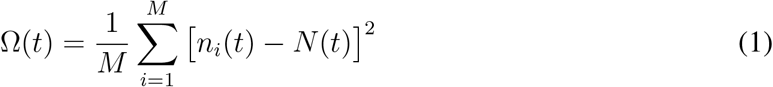

where *M* is the number of replicate simulations, *n*(*t*) is the number of micelles present in a single replicate at time *t*, and *N* (*t*) is the mean number of micelles across the replicates at time *t*. The process of the spontaneous self-assembly of micelles is considered to be sufficiently converged when Ω(*t*) is close to zero.

When determining the value of 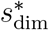 with CG simulations, it is important to also determine the corresponding radius of gyration (*R*_*g*_). There are two related observations that motivate this analysis: (1) when the CG micelle is converted to an atomistic representation with the same number of surfactants, the atomistic *R*_*g*_ is significantly smaller, and (2) the aggregation number of the atomistic micelle with the same *R*_*g*_ of the CG micelle is in good agreement with experimental DPC micelle aggregation numbers.^9,15^ The radius of gyration is given by,^20^

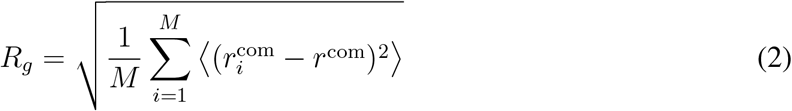

where *M* is the total number of surfactant monomers in the micelle, 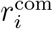 is the center of mass of monomer *i*, and *r*_com_ is the center of mass of the micelle. In order to compute the radius of gyration of the MARTINI micelles, we consider each bead to have the same mass, and therefore eq 2 simplifies to,

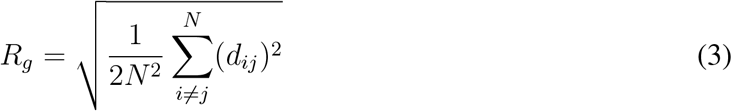

where *N* is the total number of beads in the micelle, and *d*_*ij*_ is the distance between beads *i* and *j*. The MARTINI C99 dimer was built using CHARMM-GUI^21–23^ MARTINI maker^24^ from PDB code 2LOH.^1^ The systems were packed randomly for 9 replicate simulations using the insert-molecules function of GROMACS.^25,26^ Repulsive and attractive flat-bottom harmonic biases were applied through plumed-2.8.0^27–29^ to ensure that the dimers would not aggregate together reorganize to an anti-parallel configuration. The central glycine bead (G38) of each monomer was repelled from those of every other dimer using the repulsive harmonic “lower walls” pair-distance restraint beginning at 40 Å with a force constant of 1.20 kcal/mol Å^2^. Additionally, the three intradimer distances between each G29, each G37, and each L52 were restrained with the attractive harmonic “upper walls” pair-distance restraint beginning at 17.5, 12, and 11 Å, respectively, each with a force constant of 2.40 kcal/mol Å^2^. The 9 replicates were simulated for a total of 4 *μ*s each. For the analysis, frames from every 10 ns of simulation time were used.

The MD was run with GROMACS 2020.6 with Plumed 2.8.0 on GPU.^25–29^ Dodecahedral periodic boundary conditions were used. All of the MARTINI systems were energy minimized, equilibrated, and performed at constant NPT. The systems were energy minimized using steepest descent. For MD, the leapfrog integrator was used with a 20 fs time step. The Verlet cutoff scheme for neighbor searching was applied. For nonbonded interactions, particle mesh Ewald (PME) electrostatics was applied over a range of 0-1.1 nm with an “epsilon-r” relative dielectric constant of 2.5 for all systems except for W, which had a dielectric constant of 15. For Lennard-Jones interactions, a shifting function was applied from 0.9 nm to the cutoff at 1.1 nm. For the thermostat, velocity-rescaling was used with a coupling time of 1 ps at 318 K. Isotropic Parrinello-Rahman pressure coupling was used at 1 bar with a coupling time of 12 ps and 4.5 *×* 10^−5^ bar^−1^ compressibility.

The trajectory analysis was conducted with the MDAnalysis Python library.^30,31^ The snapshots were rendered with Visual Molecular Dynamics (VMD).^32^

### Atomistic simulations of micelle-encapsulated C99

To assess the importance of choosing an accurate 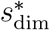 for modeling the dimeric structure of C99, atomistic DPC micelle-encapsulated dimers with two distinct aggregation numbers were built, *s*^∗^ = 100 and *s*^∗^ = 54. The 54 DPC system was chosen as a reference point based on the previously assumed aggregation number due to experimental evidence of this value for empty mi-celles.^7^ The 100 DPC system is based on the atomistic 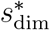 that corresponds closely to the C99 dimer-encapsulated micelle *R*_*g*_ observed in CG simulations 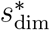. This aggregation number was ascertained by building and simulating three systems with *N* = 85, 100, and 115 surfactants, and choosing the system size that best reproduces the *R*_*g*_ from the CG model (data not shown).

The comparison between the two micelle-encapsulated dimers proceeded with the calculation of established metrics for evaluating structural properties of the C99 dimer.^7,33^ In particular, we consider *R*_*g*_,^20^ the intradimer distances between K54 and K28, the intradimer distances between all residues, the Crick angles involving G33, an intradimer dihedral angle and G33 distance,^7^ number density profiles, hydrogen bond analysis, and a measure of fractional helicity.^33^

The micelles were preassembled around the dimer using CHARMM-GUI Micelle Maker,^34^ with 5 unique initial configurations generated for each system size. The *s*^∗^ = 100 and *s*^∗^ = 54 systems were constructed in a dodecahedral perodic boundary condition and solvated with 37,000 water molecules, 8 sodium ions, and 18 chloride ions, using GROMACS insert-molecules. The initial structures of the dimer for each replicate simulation were taken from publicly available structures from previous simulation work on C99 in a POPC membrane,^35^ and processed under simulated acidic conditions such that all lysine residues were protonated.

Each replicate was minimized using the steepest descent minimization algorithm for 5000 steps with position restraints on the C99 and DPC atoms. Then, annealing was performed to thermalize the system for 2 ns, with position restraints on the same atoms as minimization. Each replicate configuration was then simulated for 1 μs, for a total of 5.0 μs of sampling per system size.

The systems were represented by the CHARMM36 protein and lipid force fields.^36,37^ The water molecules were represented by the TIP3P model.^38^ The Lennard-Jones epsilon parameter for each protein and lipid atom pair were scaled down by a factor of 0.9, based on the findings of Domanski et al. for more accurately modeling inter-peptide interactions in a lipid environment^39^ such that aliphatic hydrogen bonds can form in the C99 dimer.^33^ For the MD, GROMACS 2020.6 on GPU was used, with the leapfrog integrator with a 2 fs time step. The “Verlet” cutoff scheme for neighbor searching was applied with updates every 20 steps. For nonbonded interactions, particle mesh Ewald (PME) electrostatics from 0 to 1.2 nm and “Shift” Lennard-Jones van der Waals from 1.0 to 1.2 nm were used. For the thermostat, velocity-rescaling was used with a coupling time of 1 ps at 318 K. Isotropic Parrinello-Rahman pressure coupling was used at 1 bar with a coupling time of 2 ps and 4.5 *×* 10^−5^ bar^−1^ compressibility.

The trajectory analysis was conducted with the MDAnalysis Python library,^30,31^ with the exception of the radial distribution functions, which were computed with GROMACS. The snapshots were rendered with VMD.^32^ The set of atomistic and CG simulations performed is tabulated in Table 1.

**Table 1:** Micelle-encapsulated C99 computational assays.

| System | force field | DPC # | [DPC], M | dimer # | replicate # | sampling/replicate (μs) |
| --- | --- | --- | --- | --- | --- | --- |
| $N = 500$ | MARTINI2.2 | 500 | 0.064 | 5 | 9 | 4.0 |
| $s^* = 100$ | CHARMM36 | 100 | 0.15 | 1 | 5 | 1.0 |
| $s^* = 54$ | CHARMM36 | 54 | 0.081 | 1 | 5 | 1.0 |

## Results

### Preferred aggregation number of the C99 dimer in MARTINI

To obtain the best computationally feasible estimate of the preferred aggregation number of DPC for the C99 dimer, 5 C99 dimers and 500 DPC monomers were placed randomly in a simulation box. In the C99 dimer NMR experiments by Nadezhdin et al., the peptide concentration was reported as 0.0009 or 0.0015 M, depending on labeling method, with 50 moles of DPC per mole of peptide.^1^ Therefore, if we begin with 5 dimers, we maintain the correct ratio by including 500 DPC molecules. Further, the systems were solvated with sufficient water to dilute the DPC to a concentration of 0.064 M, and the peptides to 0.00128 M. This setup was designed to determine the unbiased aggregation number and to avoid the finite-size effects of spontaneous micellization simulations.

The evolution of the system over time is captured in Figure 1. Initially the dimers and DPC monomers are scattered randomly. After a sufficient equilibration time, an ensemble of stable micelles form, both with and without dimers inside. The ergodic measures (eq 1) of the empty micelles and all micelles are plotted in Figure 2C. The dimer micelles will always have an ergodic measure of zero (there are always 5 of them). Based on the convergence of the ergodic measure, we consider equilibration to have occurred by 2.5 *μ*s. Because of the large number of monomers and the anticipated aggregation number of around 44 for large systems of MARTINI DPC micelles based on previous work,^15^ it was expected that both dimer and empty micelles would form in sufficient quantity to avoid finite-size effects.

**Figure 1.**
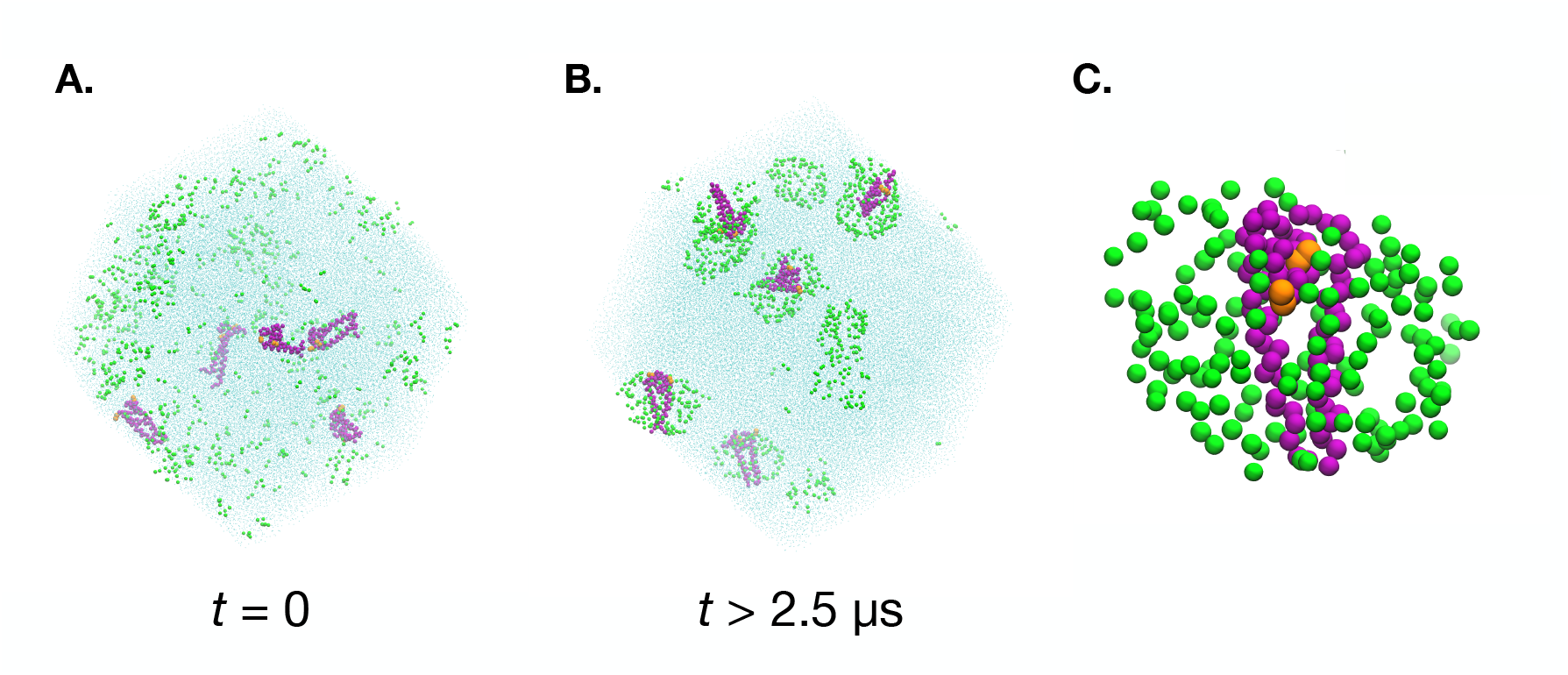
Simulation box of 500 DPC and 5 C99 dimers represented by MARTINI beads (**A**. and **B**.). The evolution of the system from a random initial configuration to an ensemble of stable micelle-encapsulated dimers and empty micelles is shown to occur over 2.5 *μ*s. A typical micelle-encapsulated dimer with the DPC headgroup beads shown in green and protein backbone beads shown in purple. The orange beads are the K28 residues (**C**.).

**Figure 2.**
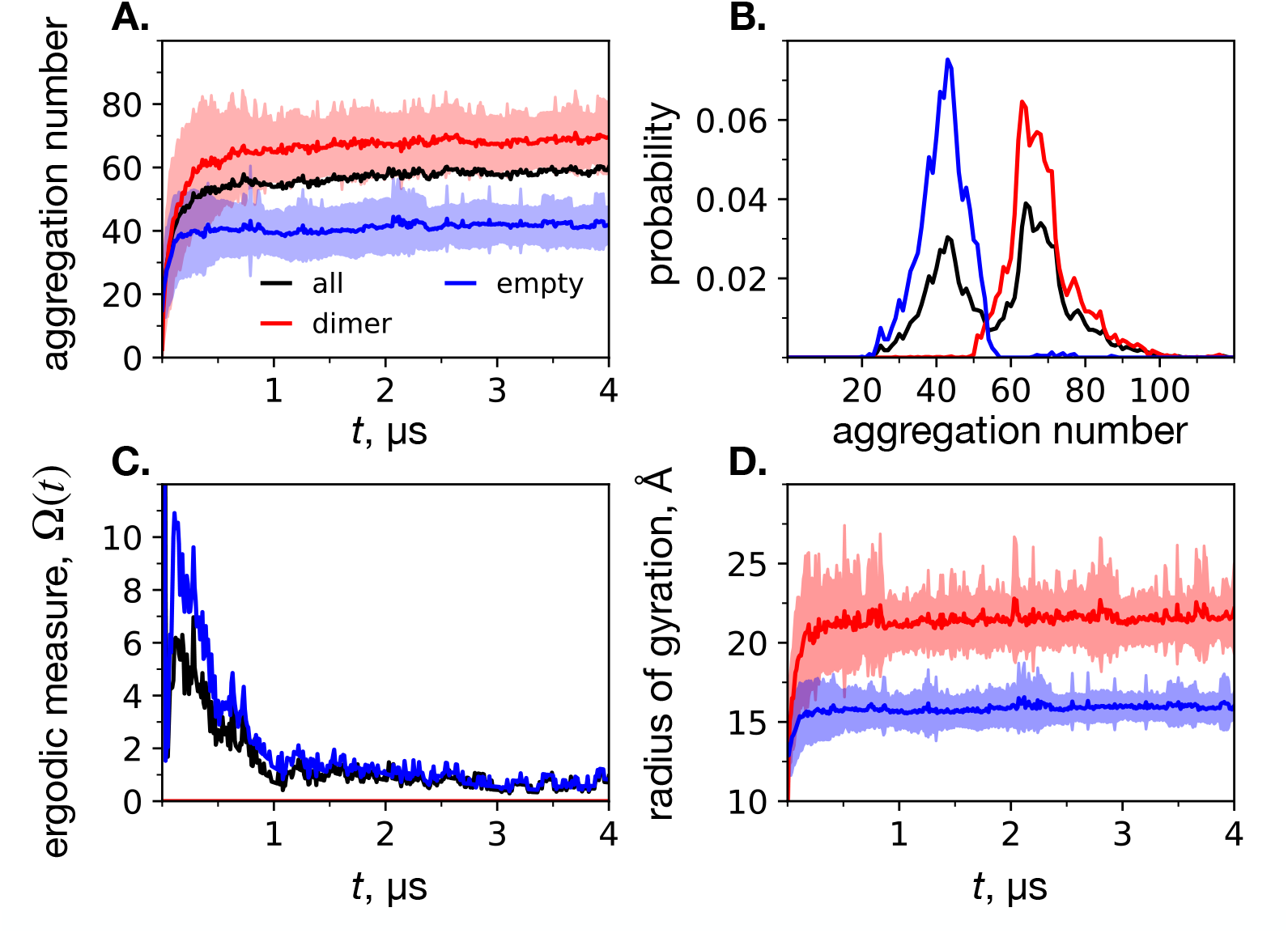
Results of CG spontaneous micellization simulations, based on three classifications of micelles: encapsulated dimer (red), empty (blue), and all (black). **A**. Mean aggregation number over time, where a minimum micelle size was defined to be 10 monomers. The error bars are the standard deviation between the replicates. **B**. Probability distribution of aggregation numbers for each of the micelle classifications. **C**. Ergodic measure (eq 1) of the convergence the systems to a stable number of empty (blue) and total (black) micelles. **D**. Radius of gyration (eq 3) over time of the encapsulated (red) and empty (blue) classifications, where the error bars are the standard deviation between replicates.

The ensemble average of the aggregation numbers of each type of micelle are depicted in Figure 2A. The average aggregation number of the second half of the trajectory (t *>* 2.5 *μ*s) is about 69 for dimer micelles, about 42 for empty micelles, and about 59 overall. Histograms of the aggregation numbers of each type of micelle are shown in Figure 2B. From the ensemble average aggregation number, as well as the histograms, it is clear that there are well-defined distributions of two distinct types of micelles. The histogram of both types of micelles is visibly bimodal, while the histograms of the individual micelles are each unimodal. Notably, the aggregation number of the empty micelles are in line with what we found previously for simulations of pure DPC with the MARTINI representation.^15^ This finding supports our assertion that the system is free from finite-size effects. In addition, this set of simulations was able to clearly show that the preferred aggregation number is significantly increased in the presence of a dimer. In other words, our simulations provide evidence that 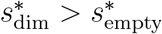, and simulations of micelle-encapsulated proteins should be performed using a surfactant number reflecting this.

The ensemble average of the radius of gyration of each micelle type over time is shown in Figure 2D. The dimer-encapsulating micelles exhibit a significantly larger *R*_*g*_ than empty micelles. The equilibrium ensemble average *R*_*g*_ is 21.6 ± 2.0 Å for the dimer micelles, and 16.0 ± 1.0 Å for the empty micelles, or about 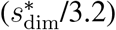 and 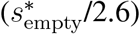, respectively. We constructed and investigated all-atom DPC-encapsulated C99 dimer systems for *s*^∗^ = 100 and *s*^∗^ = 54 DPC surfactants in the next section.

### Comparison of micelle-encapsulated dimeric structure for *s*^∗^ = 100 and *s*^∗^ = 54 DPC

To begin the comparison of dimeric structure of micelle-encapsulated C99 for two distinct aggregation numbers, we first ensure that the stability of the *R*_*g*_ value over the duration of the simulations is maintained. In Figure 3, it can be seen that each system size retains the same ensemble average of *R*_*g*_ over the entire 1 μs trajectory. For the following analysis, only simulation frames from the second half of each trajectory (*t>* 500 ns) are considered, to allow for equilibration. The mean and standard deviation in *R*_*g*_ over time and replicates is 21.83 ± 0.20 Å for *s*^∗^ = 100 and 18.21 ± 0.34 Å for *s*^∗^ = 54. Importantly, the *s*^∗^ = 100 result is consistent with the CG equilibrium dimer micelle result, and the *s*^∗^ = 54 result is consistent with the CG equilibrium empty micelle result, plus several Å due to the presence of the dimer.

**Figure 3.**
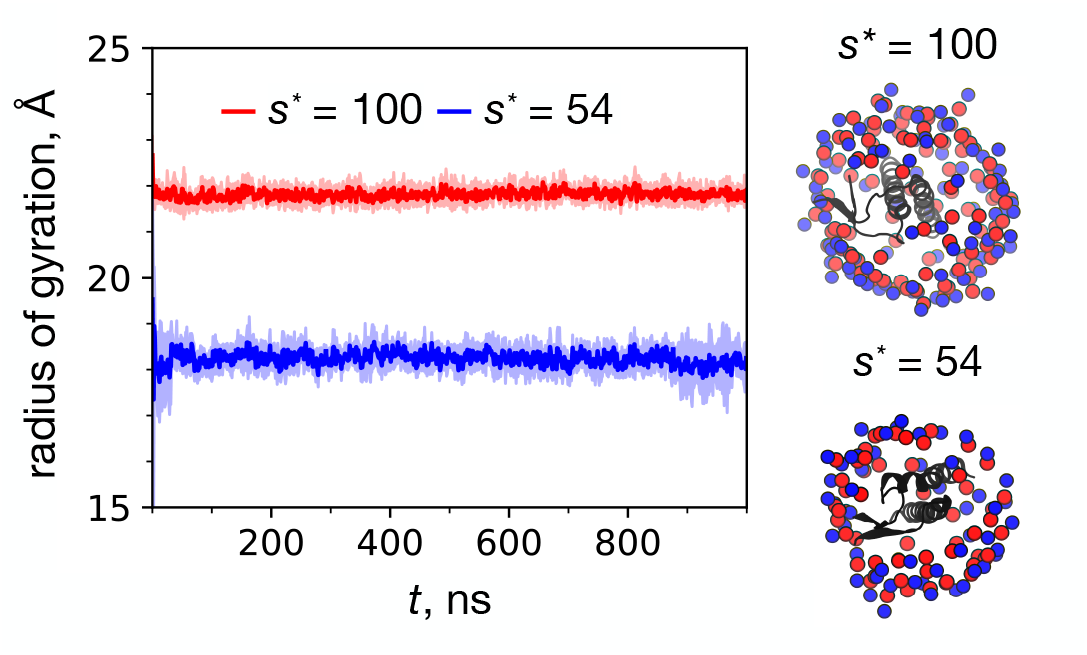
Time series of the mean radius of gyration (eq 2) for each atomistic micelle system size. The mean and error bars are based on the mean and standard deviation of the 5 replicate simulations at each time point, respectively. Also shown are representative structures of each atomistic micelle-encapsulated dimer system, viewed from above (looking down the each dimer’s helices). The two snapshots were taken with *t>* 950 ns. The DPC nitrogen atoms are blue spheres, the phosphorus atoms are red spheres, and the dimers are black ribbons.

We also note that qualitatively, the micellar structure of of the *s*^∗^=100 system is in general more stable than that of the *s*^∗^ = 54 system. This observation is supported by the snapshots of representative micelle-encapsulated dimers in Figure 3. For the *s*^∗^ = 54 system, it is apparent that the number of surfactants is not sufficient to fully cover the dimer alpha helices.

The differences in dimeric structure are first assessed through ensemble-averaged residue center of mass pair distances between all residue pairs in the dimer (Figure 4). Qualitatively, there are differences between the two distance maps, which suggest a significant impact of the aggregation number on dimeric structure. Of particular interest are the two sets of distances between the pairs of K28 and K54 residues. As indicated in Figure 4, the ensemble-averaged pair distances for K28 and K54 are 13.0 and 22.3 Å for *s*^∗^ = 100, and 15.4 and 19.8 Å for *s*^∗^ = 54, respectively. Following the analysis of Dominguez et al.^7^ for comparing the structure of the C99 dimer in membrane and micelle, we constructed a two-dimensional histogram of these sets of distances for each system size (Figure 5). The two-dimensional histograms of K28 to K28 and K54 to K54 pair distances reveals distinct conformations unique to each system size. Representative structures from simulation snapshots are shown in the bottom panels of Figure 5, indicating a tendency of the *s*^∗^ = 100 system to exhibit parallel dimers, while the *s*^∗^ = 54 assumes a crossed configuration. This observation is consistent with that of Dominguez et al. for membrane (parallel) and micelle (crossed) dimeric structures.

**Figure 4.**
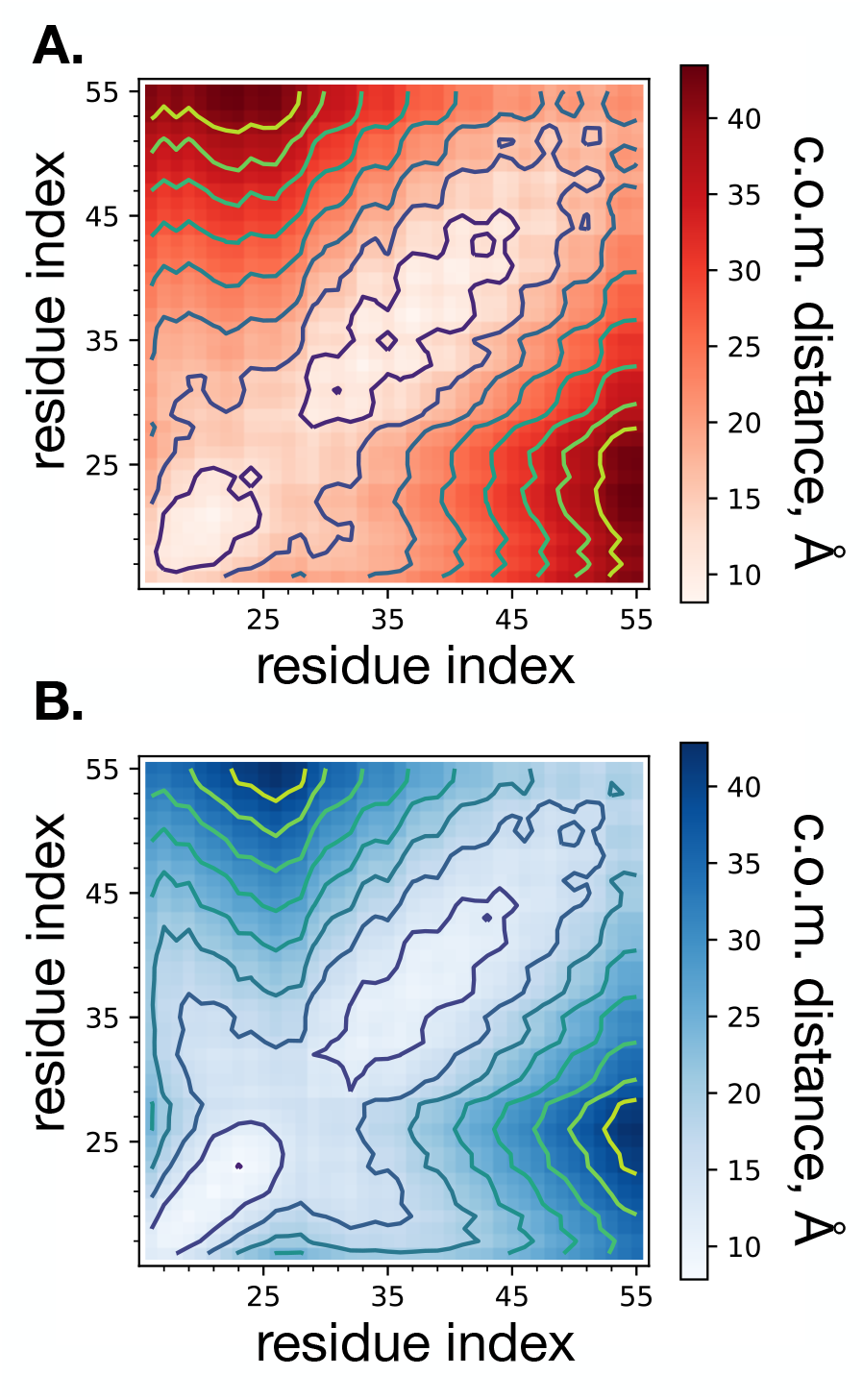
Two-dimensional color-coded image of ensemble average of center of mass distances between all residue pairs of the homodimer for each atomistic micelle system of size *s*^∗^=100 (**A**.) and *s*^∗^=54 (**B**.) surfactants. The distances are symmetrized over the diagonal, effectively averaging over the two peptides of the homodimer. There are 10 levels of distances indicated by the contour lines.

**Figure 5.**
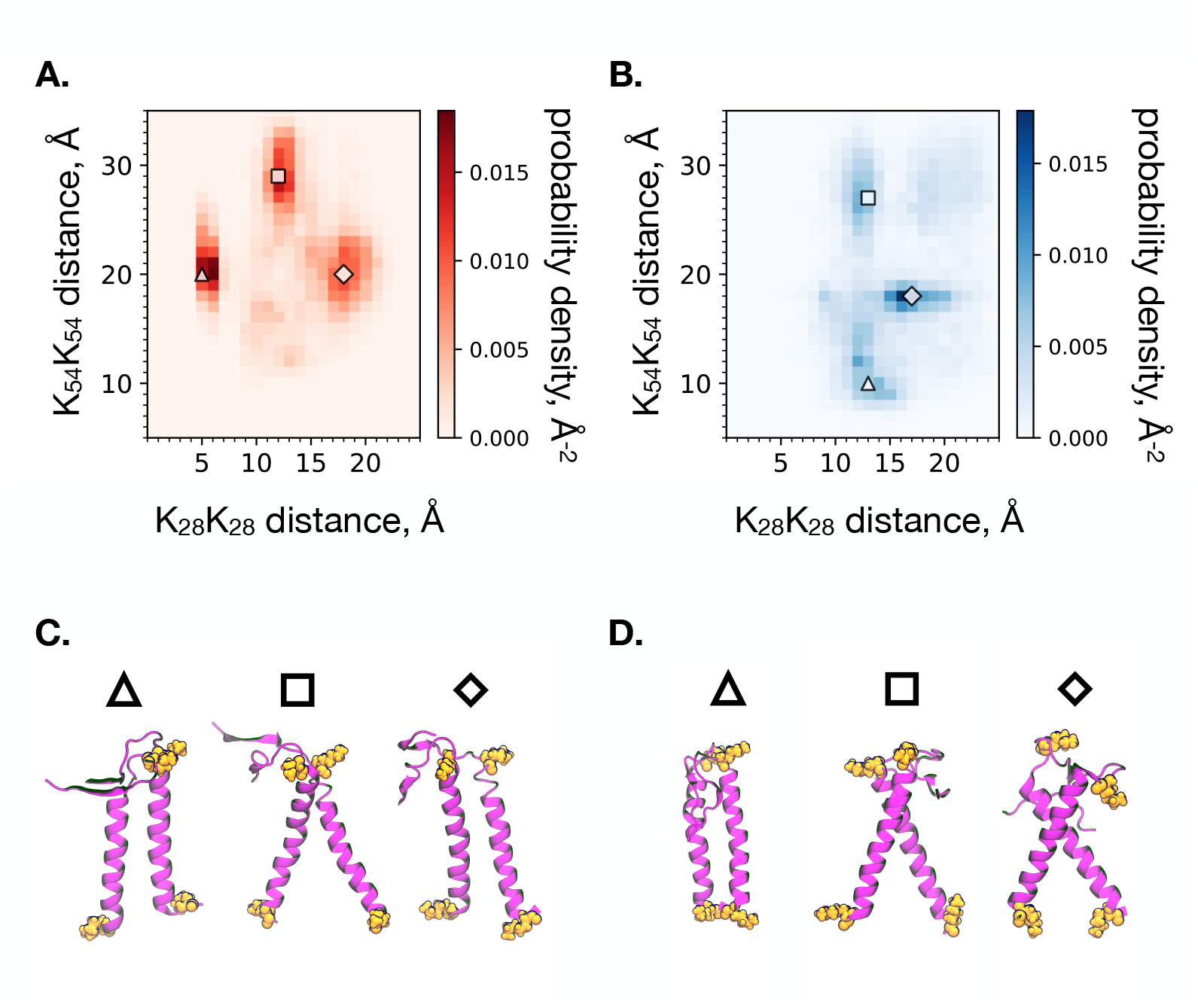
Two-dimensional histograms of the center of mass distances between the K54-K54 and K28-K28 pairs of residues for each atomistic micelle systems of *s*^∗^ = 100 (**A**.) and *s*^∗^ = 54 (**B**.) surfactants. Snapshots of representative dimer structures which correspond to the distances on the histograms marked with the corresponding shape for the *s*^∗^ = 100 (**C**.) and *s*^∗^ = 54 (**D**.) atomistic micelles. The residues K28 and K54 are colored orange (with K28 on top, K54 on the bottom), and the protein backbone is shown in purple.

To continue the comparison, we present the order parameter of the Crick angle, which is defined as the angle between the the vector connecting the two centers of mass of the two alpha helices, and the vector pointing to each G33 residue, at a normal angle to the respective alpha helix.^7^ The Crick angle describes which way the G33 residue on each peptide is pointing, with respect to the center of the dimer, ranging from 0 (“Gly-in”) to 180 (“Gly-out”) degrees. In the absence of the N-terminal extracellular domain of C99, the C99 dimer is generally stabilized by the Gly-in state, dimerizing with the same type of glycine zipper motif involving GxxxG-repeats as in the model GpA dimer.^2,33^ Two-dimensional histograms of the Crick angles of G33 for each system size are shown in Figure 6.A. (*s*^∗^ = 100) and Figure 6.B. (*s*^∗^ = 54). The distributions are similar for each system size, with both tending to adopt the Gly-in state. Dominguez et al. had reported that the Gly-in state is more likely in lipid bilayer, and had observed that Gly-out was more likely to occur in micelle.^7^ As may be expected from increasing the curvature of the micelle surface, we see there is a higher likelihood for the Gly-out state in *s*^∗^ = 54 compared to *s*^∗^ = 100 micelles, which can be seen in the population in the 135-180 degree region of Figure 6.B.

**Figure 6.**
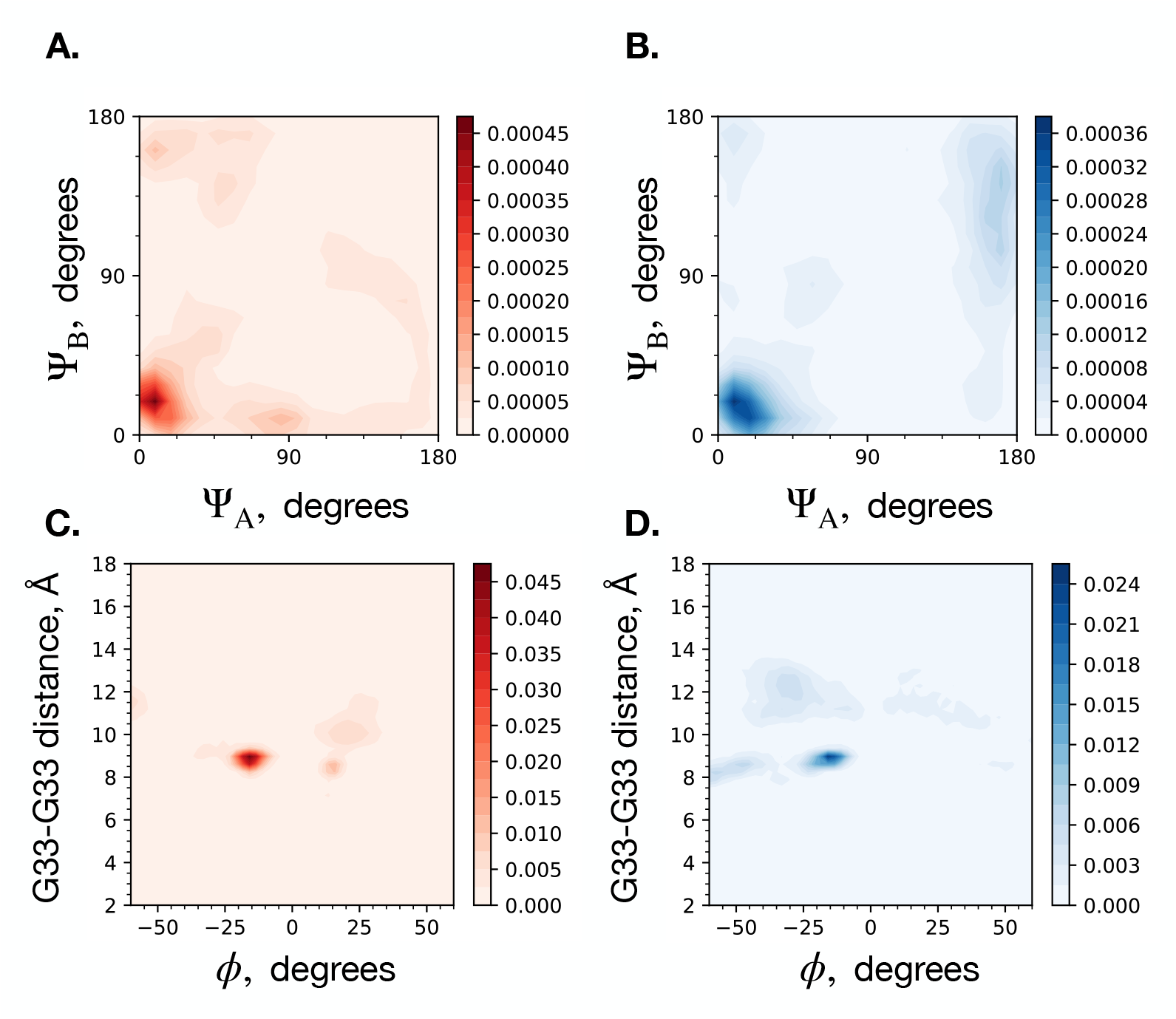
Two-dimensional histogram of crick angles of G33 on each C99 congener (Ψ_*A*_, Ψ_*B*_) within the dimer for the *s*^∗^ = 100 (**A**.) and *s*^∗^ = 54 (**B**.) atomistic micelle systems. ^33^ Two-dimensional histogram of G33-G33 alpha carbon pair distance and dimer twist *ϕ* for the *s*^∗^ = 100 (**C**.) and *s*^∗^ = 54 (**D**.) atomistic micelle system sizes. ^33,40^ The colorbars correspond to the normalized ensemble probability densities.

Another meaningful order parameter is the dihedral angle, *ϕ*, of G29A-G37A-G37B-G29B, where A and B refer to peptide A and peptide B of the C99 dimer.^7^ The sign of *ϕ* indicates the handedness of the coiled-coil of the dimeric structure. A negative value of *ϕ* corresponds to a right-handed coiled-coil, and a positive value corresponds to a left-handed coiled-coil. To further compare the dimeric structure between system sizes, we constructed a two-dimensional histogram of *ϕ* and the intradimeric distance of the pair of G33 residues. These distributions are shown in Figure 6.C. (*s*^∗^ = 100) and Figure 6.D. (*s*^∗^ = 54). These are also similar between the two system sizes. The predominantly negative (right-handed) *ϕ* value is consistent with what was found by Dominguez et al. for the C99 dimer in both membrane and micelle.^7^

To further elucidate the dimeric structure in each micelle, we analyzed the propensity for backbone hydrogen bond formation between all pairs of residues. The glycine zipper of the transmembrane domain is expected to facilitate dimerization of the transmembrane domain via formation of the Gly-in state, stabilized by aliphatic hydrogen to carbonyl oxygen hydrogen bonds (C_*α*_*H ···* O=C) involving G29, G33, G37. Likewise, the juxtamembrane domain residues 16-28 are expected to potentially form inter-C99 beta-strands (NH*···* O=C) if not at a pH or lipid phase which facilitates helix formation in residues 16-23.^2,33^ Using the ensemble-average propensities for hydrogen bonds between pairs of residues, *i* and *j, P*_*ij*_, we computed two-dimensional and one-dimensional probability distributions for each of the two types of hydrogen bonds and for each micelle size (Figure 7). In micelles of size *s*^∗^ = 100, we find that the aliphatic hydrogen bonds characteristic of the glycine zipper, which stabilize the Gly-in state, form with relatively greater propensity than in *s*^∗^ = 54 micelles, particularly at G29 (Figure 7.A,B,C,F). Parallel beta strands involving residues 16-24 sometimes form in *s*^∗^ = 54, and can bury into the micelle and interact with the transmembrane domain (Figure 7.E), which does not happen in *s*^∗^ = 100. These parallel beta strands between juxtamembrane domains in *s*^∗^ = 54 micelles form when the micelle has transient defects, which we show in the following section allow water to penetrate into the micelle. In *s*^∗^ = 100 micelles, the juxtamembrane domains of C99 form transient interactions with each other, but not extended beta strands (Figure 7.D).

**Figure 7.**
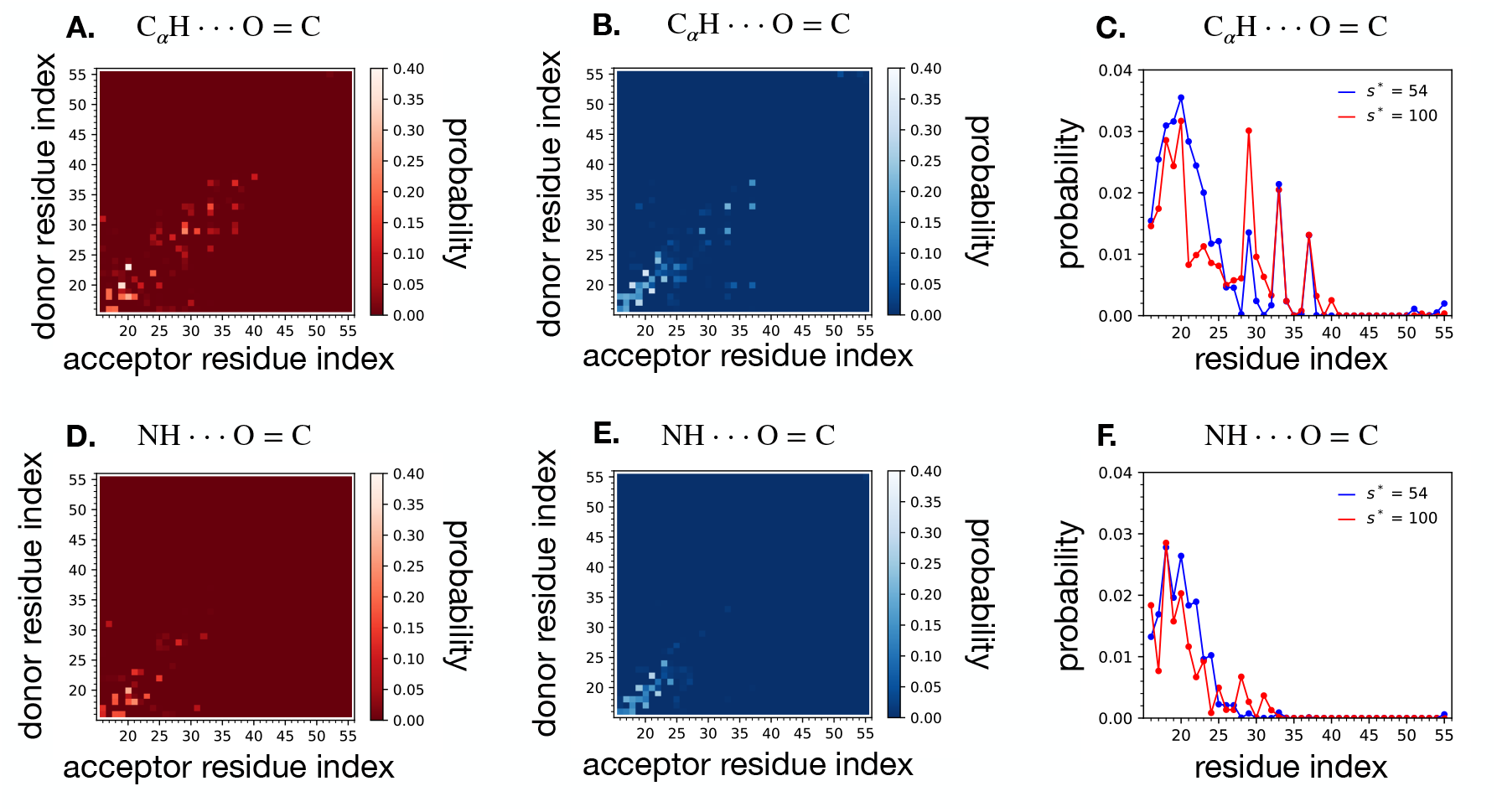
Two-dimensional histogram of intradimer hydrogen bond probability between each pair of alpha carbon hydrogen and backbone carbonyl for the *s*^∗^ = 100 (**A**.) and *s*^∗^ = 54 (**B**.) atomistic micelle systems. A hydrogen bond is counted if the relevant atoms are within 3.5 Å and form an angle of at least 120 degrees. The probability is normalized by the total number of simulation frames included in the calculation. The one-dimensional probability distribution of participating in the hydrogen bond (**C**.) for each residue *i* was computed as the mean of column *i* plus the mean of row *i* in the corresponding two-dimensional histogram. Bottom row: Two-dimensional histogram of intradimer hydrogen bond probability between each pair of backbone nitrogen hydrogen and backbone carbonyl for the *s*^∗^ = 100 (**D**.) and *s*^∗^ = 54 (**E**.) atomistic micelle systems, and corresponding one-dimensional probability distributions (**F**.). These hydrogen bonds are counted if the relevant atoms are within 2.5 Å and form an angle of at least 150 degrees.

The structure of each peptide within the dimer in micelles of each aggregation number was also evaluated through the calculation of fractional helicity. The fractional helicity value is the observed propensity for each residue to be in an alpha helix. It is defined for each residue, *i*, by the ensemble-averaged number of backbone hydrogen bonds between residue *i* and *i* ± 3, 4, 5. hydrogen bonds must be within 2.5 Å at an angle of at least 150 degrees. The fractional helicity (Figure 8) of C99 was found to be similar in *s*^∗^ = 100 and 54 in the transmembrane domain, but different in the juxtamembrane domain. In the juxtamembrane domain, the fractional helicity of C99 in *s*^∗^ = 100 micelles in higher in residues 20-25, attributable to the beta strand that can form with these residues in *s*^∗^ = 54 micelles.

**Figure 8.**
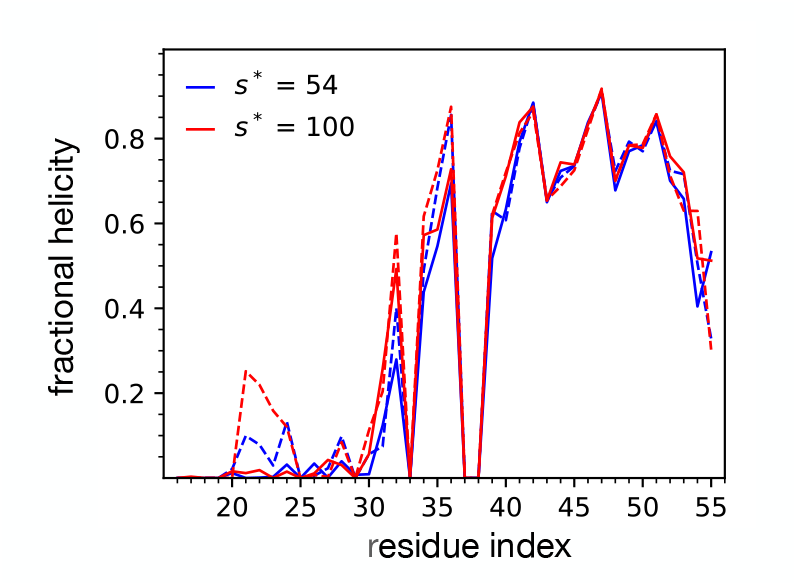
Fractional helicity order parameter for each atomistic micelle-encapsulated C99 dimer. Fractional helicity is the probability that a residue is participating in a peptide backbone hydrogen bond, which is defined for each residue, *i* by the ensemble-averaged number of backbone hydrogen bonds between residue *i* and *i* ± 3, 4, 5. The plotted lines correspond to the calculated values for peptide A (solid line) and peptide B (dashed line), for the *s*^∗^ = 100 (red) and *s*^∗^ = 54 (blue) system sizes.

Aside from dimeric structure, we also conducted a systematic analysis of the atomistic micellar structure in terms of the radial distribution function (RDF). We calculated the number density, or RDF without density normalization, as implemented in the GROMACS RDF function^26^ for several structural features of each micelle-encapsulated dimer system. In particular, number density RDFs of DPC surfactant nitrogen, DPC carbon, water oxygen, and protein alpha carbon atoms are provided in Figure 9. The number densities for each system size are plotted together and underscore the significant size-dependent differences in micellar structure. In each case, *s*^∗^ = 100 is the solid line, and *s*^∗^ = 54 is the dashed line. For consistency in the comparison of the structure of each micelle, the same reference (or origin) position is used for each density profile, which was chosen to be G33 of peptide A. The DPC nitrogen number density profile clearly demonstrates the increased micellar radius of the *s*^∗^ = 100 system, with a larger number density further from the reference point. The DPC carbon number density profile follows the same observation as that of DPC nitrogen. In the water oxygen profiles, the number density far away from the reference reaches a plateau of around 0.033 water oxygen per Å^2^ for both system sizes. This value corresponds to a concentration of 55 M, close to the expected concentration of water (and allows us to double check our units: (0.033 Å ^−3^*/N*_*a*_) *×* (10^24^ Å^2^*/*cm^3^) *×* (1000 cm^3^*/*L) = 55 mol*/*L). For the short distance region, however, the number density of water is higher for the smaller micelle, indicating that the dimer is more exposed to water. We interpret the water density peak occurring at around 5 Å as the extent of water penetration for each micellar system. The larger water peak for the *s*^∗^ = 54 system can also be partially accounted for by micellar deformation such as that captured in Figure 3. Lastly, the number density profiles of the dimer alpha carbon atoms are similar for each system size. An exception is around the distance of 13 Å, where the *s*^∗^ = 54 system has a slightly higher density than the *s*^∗^ = 100 system.

**Figure 9.**
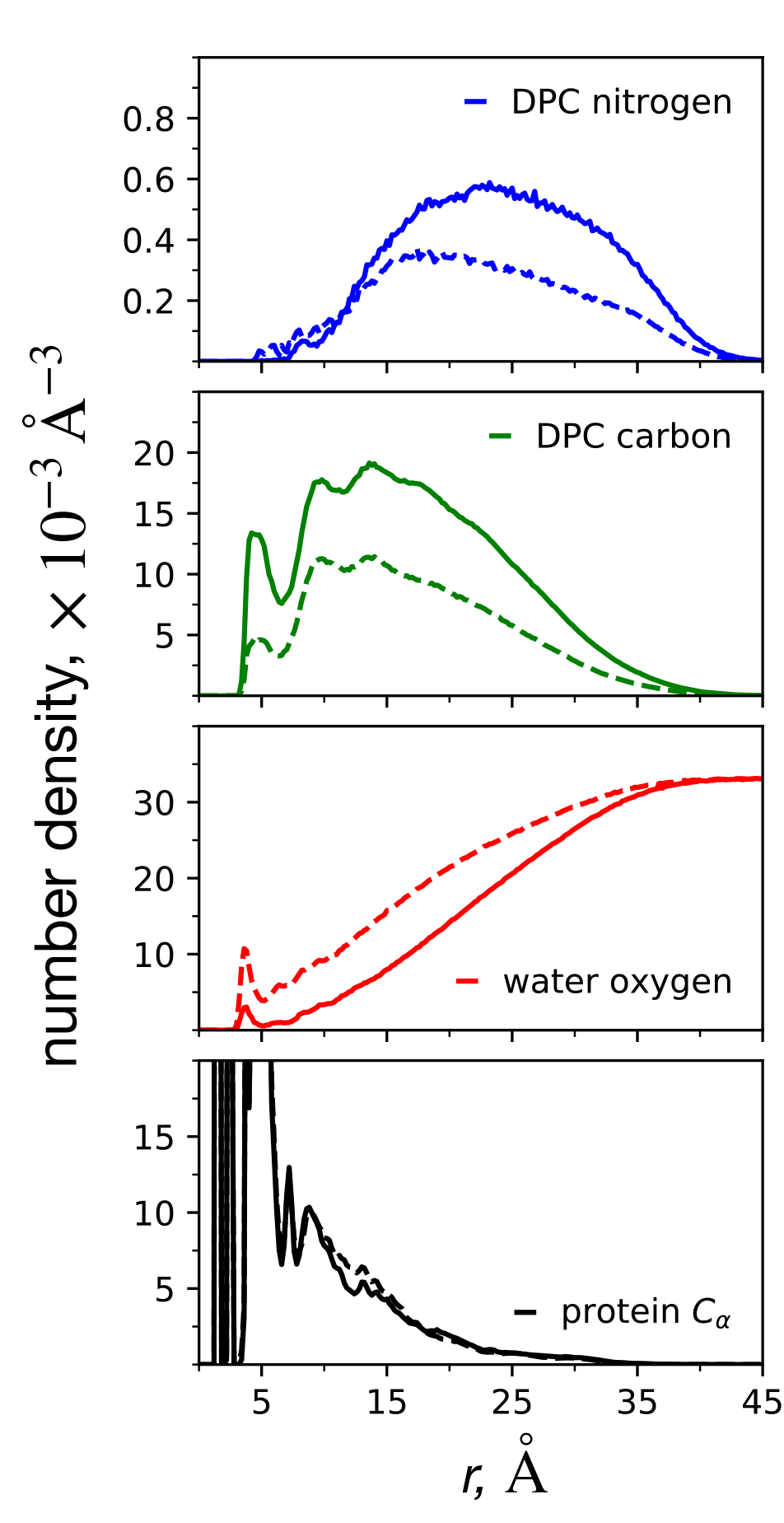
Number density radial distribution functions describing the C99-micelle encapsulated structure for the *s*^∗^ = 100 (solid lines) and *s*^∗^ = 54 (dashed lines) atomistic micelle systems. In each case, the reference position is the alpha carbon of Gly33 peptide A. Separate number density profiles are computed for the head-group nitrogen atoms (blue), each DPC carbon atom (green), the oxygen of every water (red), and the protein backbone atoms (black).

## Conclusion

Through the multiscale modeling of the DPC micelle-encapsulated C99 dimer presented in this study, we demonstrated (1) that the preferred aggregation number of a DPC micelle around the dimer is significantly larger than that of the empty DPC micelle and (2) that the transmembrane structure of the C99 dimer depends on the aggregation number of its micelle. By conducting similar analysis on the transmembrane dimeric structure to that of previous studies of the dimer in micellar and membrane environments, we provide evidence that the dimeric structure in the larger, more accurate micelle is more similar to the dimer in the membrane environment than the smaller micelle. Additionally, the smaller micelle was observed to have defects in which the dimer transmembrane region is temporarily exposed to water, which is evidenced by simulation snapshots, the instability in the time series of the mean radius of gyration, and the relatively pronounced short-range water peak in the number density profile. The relatively higher propensity for hydrogen bonds in the glycine zipper in larger micelles relative to smaller micelles suggests that the C99 dimers in larger micelles are more stable, and that reducing the curvature of micelles and lipid bilayers may generally encourage C99 dimerization. These observations indicate that it can be valuable to expend the computational effort to determine the accurate preferred aggregation number in the presence of the protein, when attempting to model a transmembrane structure using a micelle as a membrane mimic.

We have shown how the preferred aggregation number can be ascertained conveniently using CG simulations, without any estimate of its value a priori, and then converted to an equivalent atomistic value. Our approach was built on the findings of our previous work on finite-size effects in simulations of micelle self-assembly, which concluded that there should be sufficient monomers for at least three micelles to form. This finding is extended to micelle-encapsulate proteins by ensuring that at least three empty and three protein micelles are able to form. While the precise minimum system size is not known beforehand, the resulting micelle distribution can be quickly analyzed to check if the finite-size criteria are met. This method was made computationally feasible by our ability to rely on CG simulations for the large self-assembly simulations, due to the known relationship between the CG radius of gyration and the accurate atomistic preferred aggregation number: while the CG aggregation number does not correspond to the accurate atomistic aggregation number, the CG radius of gyration does correspond to an accurate atomistic micelle’s radius of gyration. Atomistic representations of the micelle-encapsulated protein are therefore able to be built pre-formed based on the radius of gyration obtained from the large CG simulations.

While we conducted unbiased atomistic simulations of the C99 dimer in micellar environment with a total of 5 μs of sampling per system, it may be necessary in future work to implement an enhanced sampling strategy such as replica exchange or metadynamics in order to fully characterize the metastable states of the dimer’s conformational free energy landscape.

## Acknowledgement

The authors gratefully acknowledge the generous support of the National Science Foundation (CHE-1900416), the National Institutes of Health (R01GM107703), and the high-performance computing resources of the Boston University Shared Computing Cluster (SCC).

